# Nanoscale resolution of microbial fiber degradation in action

**DOI:** 10.1101/2021.02.16.431430

**Authors:** Meltem Tatlı, Sarah Moraïs, Omar E. Tovar-Herrera, Yannick Bomble, Edward A. Bayer, Ohad Medalia, Itzhak Mizrahi

**Affiliations:** Department of Biochemistry, University of Zurich, Zurich, CH-8052, Switzerland; Faculty of Natural Sciences, Ben-Gurion University of the Negev, Beer-Sheva 8499000 Israel; Biosciences Center, National Renewable Energy Laboratory, Golden, CO USA; Department of Biomolecular Sciences, The Weizmann Institute of Science, Rehovot 7610001 Israel

## Abstract

Deconstruction of plant cell walls is imperative to global carbon cycling and sustainability efforts. Selected microbes degrade plant fibers using extremely efficient multi-enzymatic cellulosomes assemblies. Organization of cellulosomes on the bacterial cell surface and their ecological regulation remain elusive. By combining structural methodologies with molecular and biochemical approaches on the canonical *Clostridium thermocellum* system, we provide an unprecedented view into the in-situ structure and distribution of cellulosomal enzymes while interacting with their cellulosic substrate during fiber degradation. Structural exploration of growing cultures revealed isogenic phenotypic heterogeneity of cellulosome organization on single cells across the bacterial population, suggesting a division-of labor strategy driven by product-dependent dynamics. This study demonstrates how structural biology under near-physiological conditions can be employed to develop ecological hypotheses to understand microbial plant-fiber degradation at the single-cell nanoscale level.

**One Sentence Summary:** This study contributes critical insights into the *in-situ* organization of cellulosomes and their cellulosic substrates and provides evidence for phenotypic heterogeneity, with dynamic, growth phase-dependent organization of the fiber-degrading machinery.

## Introduction

Plant biomass is the largest source of biomass on planet Earth ^1^. Hence, plant cell wall degradation is an essential and fundamental process in nature with major consequences to carbon recycling, environmental homeostasis and of major significance to waste management and mammalian gut ecosystems. Bacteria are central players in this process with the essential role of deconstructing cellulosic fibers into simple sugars that could be used by a multitude of organisms as well as in industrial processes ^2^. Several anaerobic bacteria evolved an intriguing, energy-effective strategy to dismantle plant fiber by the production of cellulosomes, i.e., multi-enzymatic complexes that evolved to be highly-efficient in fiber degradation ^3^.

Cellulosomes are massive multi-modular protein machineries, composed of multiple enzymes and enzyme-integrating subunits (“scaffoldins”) that are highly efficient in deconstructing plant cell wall polysaccharides ^4–6^. Although extensive research has been dedicated to cellulosome activity and their composition in various cellulosome-producing bacteria ^3^, an understanding of how these complexes are distributed on the bacterial cell wall and interact with their substrates is still unclear. Thus, *in-situ* structural characterization of cellulosome complexes remains a challenging task ^7^.

Major hurdles for high-resolution imaging of cellulosomes include inherent compositional heterogeneity, high flexibility of their elongated linkers and the variety of alternative conformations that reflect the dual-binding mode that characterizes the integration of the enzymatic subunits into the structural scaffoldin proteins that are serving as the scaffold on which the cellulosomal complex is assembled. Despite these obstacles, conventional methods have identified structural features of individual cellulosomal modules, as well as crucial interface residues that are responsible for the specific tenacious intermodular binding between the definitive cellulosomal modules ^8–15^. In addition, larger cellulosome fragments have been structurally studied by complementary approaches, i.e., crystallography, smallangle X-ray scattering, computational modeling and negative-staining transmission electron microscopy ^16–20^. Additional studies have revealed some of the structural features of a cellulosome fragment composed of five separate cellulosomal modules from three different proteins ^21–23^. More recently, truncated recombinant minicellulosome complexes, from the canonical and most-studied cellulosome-producing bacterium *Clostridium thermocellum*, comprising a truncated scaffoldin and three copies of the Cel8A cellulosomal endoglucanase, have been analysed ^19^. These *in-vitro* studies, focused mainly on specific enzymatic and structural components of the complex, without the presence of the microbial cells on which the cellulosomal complexes are situated. Pioneering studies published in the 1980s, using electron microscopic imaging, have provided a two-dimensional glimpse into the massive size and organization of the cellulolytic machinery on the bacterial cell surface of *C. thermocellum* ^3,4,20,24,25^. Nevertheless, these studies employed dehydration and chemical fixation procedures, which thus restrict the structural information and characterization of these bacterial cell wall systems.

Hence, robust insight into the fiber-degrading process at the nanoscale level of a bacterium, while these complexes are in the active state, remains unknown. Moreover, the question of how and whether the molecular organization of these complexes on single cells is interconnected with functional and regulatory aspects of cellulosomes at the population level is essential to our understanding of the fiber-degradation process and these canonical enzymatic complexes.

To answer these questions and circumvent the aforementioned obstacles, we exploited current technological advances that combined cryo-electron microscopy (cryo-EM) ^26^ and cryo-electron tomography (cryo-ET) ^27,28^ together with conventional approaches of microbial genetics, physiology and biochemistry. Using these complementary approaches, we gained deep insight into the *in-vitro* and *in-situ* structure of the cellulosome enzymes, from the single-enzyme level to the distribution of cellulosome complexes across single cells in a population, to reveal how cellulosome structure and distribution may be connected to gene expression and regulation at the population level.

## Results

### Cellulosomes are imaged around intact bacteria as an extensive layer of continuous complexes of constant distance from the cell wall

To examine the detailed organization of the cellulosome complexes *in situ*, we applied cryo-EM and cryo-ET imaging to intact *C. thermocellum* bacterial cells. Low-magnification images of the bacteria indicated a typical view of the elongated cells, measuring a few micrometers in length and 500-600 nm in thickness (Fig. 1A), which represents a suitable size for *in toto* analysis of bacteria by cryo-ET ^29^. Closer inspection of these bacteria confirmed the presence of a dense layer surrounding the bacterium (Fig. 1A, arrowhead). *S*lices through typical cryo-tomograms (Fig. 1B,C) revealed discrete globular protein densities in the extracellular layer and densities that emanate from the bacterial S-layer to this extracellular protein layer (Supp. Fig. S1).

**Fig. 1.**
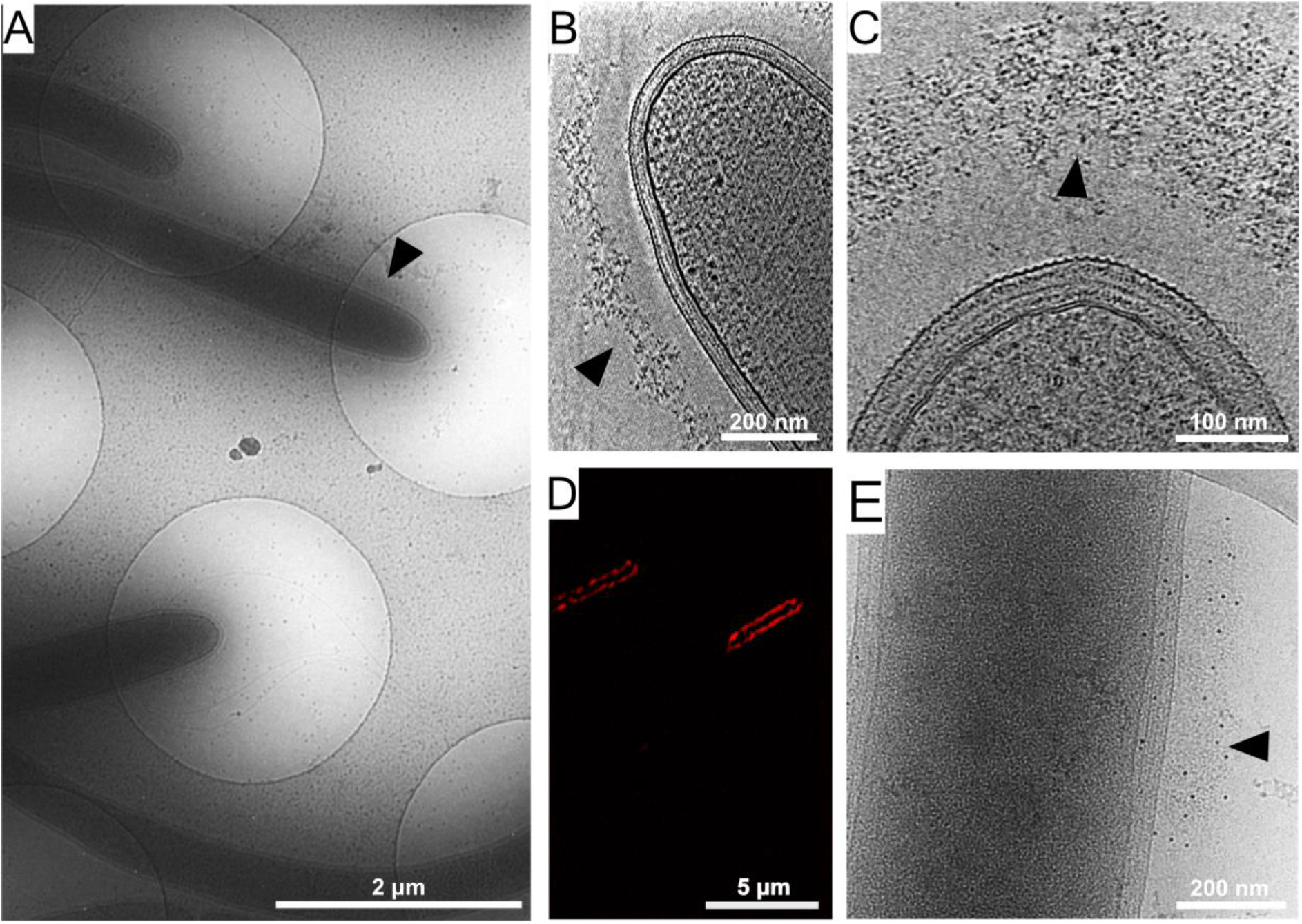
Cellulosome organization around *C. thermocellum* bacterial cells. (A) An image of *C. thermocellum* cells with surrounding cellulosomes on a cryo-EM grid at low magnification. The black arrow indicates the grey hue that forms a layer of extracellular cellulosome complexes surrounding the bacterium. At this magnification, the layer seems homogeneously thick. (B,C) Cellulosome complexes of two different bacteria at different magnifications. In (B), cellulosomes form a condensed layer which varies in thickness (arrowhead). At higher magnification in (C), the tomographic slice reveals the cellulosomes in greater detail (arrowhead). The density of the cellulosome layer reflects the protein occupancy within the layer. (D) Immunofluorescence confocal microscopy image of *C. thermocellum* bacteria reveals the position of the cellulose-binding module 3a (CBM3a) component of the CipA (ScaA) scaffoldin around the cells. (E) The extracellular density (arrowhead) detected by cryo-EM was verified to be cellulosomes. Immunogold labelling of cellulosomes, applied to localize the CipA scaffoldin using anti-CBM3a antibodies, indicates that the density detected around the bacteria corresponds to cellulosome complexes. The colloidal gold appears as black dots (arrowhead), while control experiments showed no gold nanoparticle signal around the bacteria (see Fig. S2A,B).

To confirm that the extracellular density layer detected by cryo-ET is composed of cellulosome assemblies, we utilized confocal immunofluorescence imaging and immunogold labelling with specific antibody targeting of the cellulose-binding module 3a (CBM3a) component of the CipA (ScaA) scaffoldin, the major defining constituent of the *C. thermocellum* cellulosome complex. The results of these experiments revealed a continuous layer of cellulosome structures around these bacteria (Fig. 1D), whereby gold nanoparticles were detected at the densities around the cells that correspond to cellulosome assemblies (Fig. 1E, Fig. S2B and control Fig. S2A).

To further verify that the densities detected correspond to the fiber-degradation machinery, we analysed two mutant strains of *C. thermocellum*, which either lacked the CipA scaffoldin subunit or lacked all of the anchored scaffoldins (knockout mutants DSN11 and CTN7, respectively) ^30^. Cryo-ET analysis of these mutant bacteria indicated complete absence of extracellular densities as compared to the wild-type cells (Supp. Fig. S2C-F). The above experiments suggest that the densities detected (Fig. 1) correspond to the cellulose-degradation machinery of *C. thermocellum*.

Surprisingly, cryo-tomograms of intact *C. thermocellum* cells revealed a characteristic gap between the S-layer and the cellulosome layer, with an average distance of 64±17 nm, while the thickness of the cellulosome-rich layer was 65±16 nm (Supp. Fig. S3), measured for 24-h grown bacterial cultures. However, density was also observed that emanated from the bacterium to the cellulosome layer, which presumably indicated the anchorage of the cellulosome to the cell wall (Supp. Fig. S1, arrows). The gaps are reminiscent of the previously described “contact corridors” ^24^, which were proposed to contain anchoring proteins, e.g., OlpB (ScaB), that include exceptionally lengthy intermodular linking segments ^25,31,32^.

### Bacterial cell surface-associated enzymes

The *C. thermocellum* genome contains genes that encode 72 different dockerin-containing proteins, which result in heterogeneous cellulosomal configurations. A number of proteomic studies have revealed the abundance of the different cellulosomal enzymes. Exoglucanase Cel48S has been unanimously reported as the most expressed dockerin-containing enzyme and crucial for enzymatic digestion of microcrystalline cellulose ^33–36^. We therefore studied the structure of recombinantly expressed full-length Cel48S by cryo-EM. Figure 2A shows the determined 3.4 Å resolved structure (Supp. Fig. S3). The structure highly resembles the previously reported crystallographic structure that lacks the linker and dockerin module^37^, although, in the current cryo-EM structure, an additional stretch of 10 amino-acids (MTYKVPGTPS) can be detected at the surface of the protein, which represents the major part of the linker between the catalytic module and the dockerin (Fig. 2A, purple). Curiously, the dockerin, although encoded, expressed and confirmed by mass-spectrometry, was not visible in the high-resolution density map, presumably due to the flexibility of the hinge region (TKLYG) that bridges the enzyme’s catalytic module with the dockerin, which would allow conformational freedom to interact with the substrate.

**Fig. 2.**
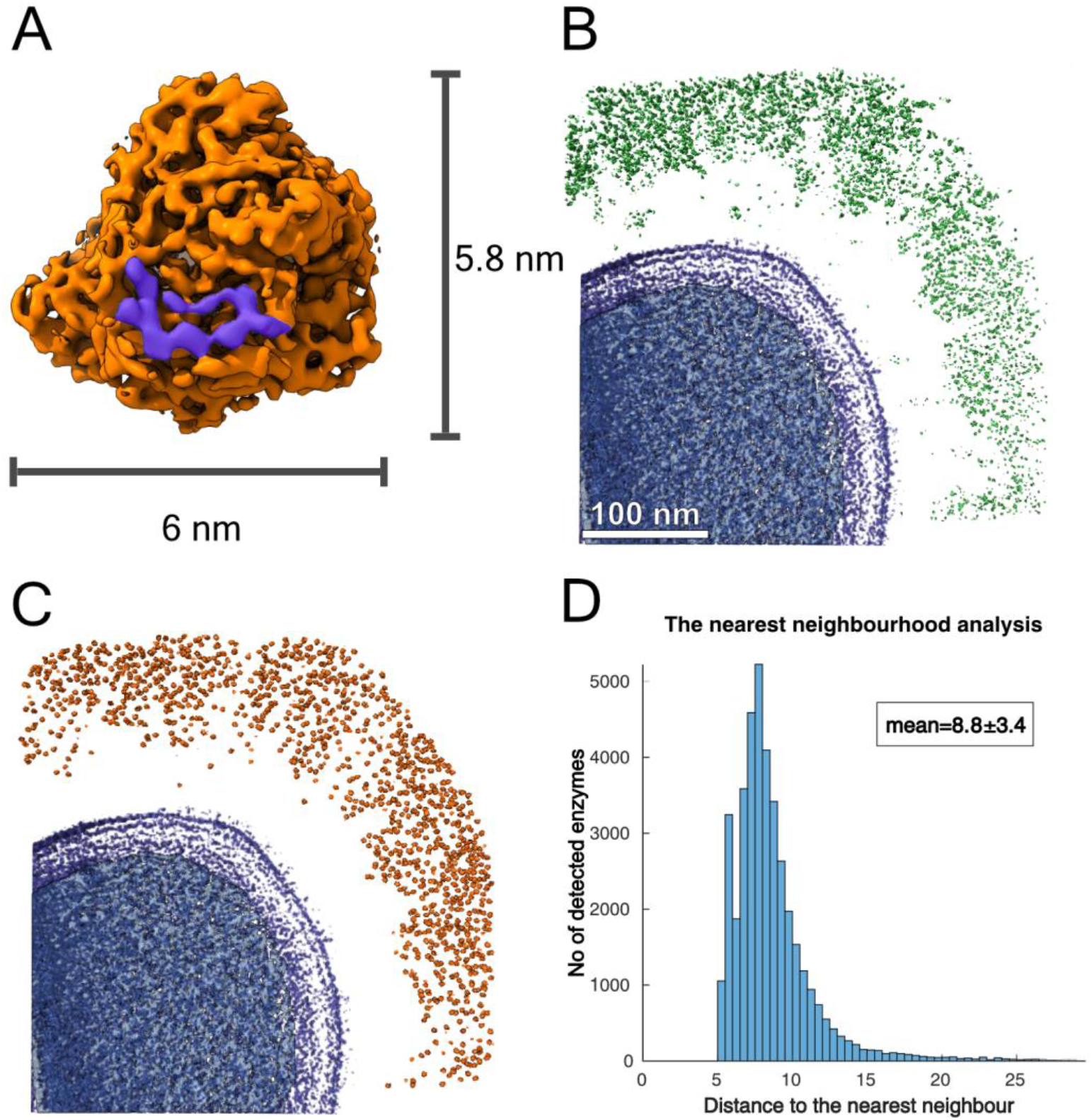
Density of cellulosomal enzymes around *C. thermocellum*. (A) High-resolution structure of the full-length Cel48S was determined by cryo-EM to a resolution of 3.4 Å (Supp. Fig. S4). The purple region is part of the linker that was structurally determined between the enzyme and the dockerin. (B) A rendered view of the cellulosome organization (green) around the bacterial cell wall (purple) depicted in central volume was used for template-matching analysis. A threshold level was applied to the entire volume. The positions of the Cel48S enzyme and similar globular densities around the bacterium were determined by template matching and displayed in (C, orange). The coordinates and angles of the enzyme were determined by means of template matching, using the structure determined by cryo-EM (A) as a template, and mapped back into the cryo-tomograms. (D) The shortest distance between neighbouring enzymes was determined and shown for 8 bacteria (tomograms in Supp. Fig. S7).

Since Cel48S is the most abundant enzyme in the cellulosome, we used the determined structure to identify enzymatic subunits within the dense cellulosome layer around the bacteria. To this end, a template-matching algorithm was applied onto cryo-tomograms of the bacteria, as previously described ^38^. Eight tomograms which exhibit different densities of cellulosome organization were depicted for this experiment (rendered views of two bacteria are shown in Fig. 2B). The matched densities of the Cel48S (Fig. 2C, orange) in the protein layer around the cells resembled the dimensions of the structure described above (Fig. 2B), suggesting that a major part of the observed densities corresponded to the Cel48S and other cellulosomal enzymes, as previously shown ^39^. Indeed, manually segmented cellulosome densities (Fig. 2C) resulted in a similar pattern as the above-described template-matching experiment, thus confirming the high abundance of the enzyme.

The distributions of distances between neighbouring enzymatic subunits around the eight bacteria are presented in Fig. 2D (analysis of individual tomograms are shown in Fig. S5). The mean distance of neighboring particles was calculated to be 8.8 nm, which resembles the distance between enzymes in reconstituted cellulosomes ^19^. These results suggest that 20 to 50% of the volume of the cellulosome layer is occupied by Cel48S subunits, which are positioned in close proximity to each other, presumably to increase the efficiency of fiber degradation. The average thickness of the cellulosome layer detected was measured at 65 ± 16 nm (Fig. S4). Thus, a maximum of 69,000±16,000 volumes of Cel48S per micrometer of the bacterium could be accommodated within the cellulosome volume that encompasses the bacterium. However, due to the spacing between the cellulolytic enzymes (Fig. 2D), 14,000-35,000 enzymatic subunits around a micrometer of the bacterium are estimated.

### Cellulosomes caught in action

How does the cellulosome interact with its cellulose substrate? Unprecedented insight into cellulosome-substrate interactions around the bacterial cell surface was obtained by utilizing volta-phase plate (VPP) assisted cryo-ET analysis, which provides high contrast imaging without the need for additional processing ^40^. We thus cultured *C. thermocellum* in the presence of microcrystalline cellulose and were able to identify interactions between cellulose and the cell-associated cellulosomes. Strikingly, we observed the cellulosomes, which form a layer around the bacterium and envelop the cellulosic fibers. The cellulose was identified within the dense cellulosome layer, suggesting flexible organization of the enzymes within the cellulosome layer while encompassing the cellulose substrate, presumably offering an advantage for its degradation (Fig. 3A). To further understand the enzyme-substrate interface and to obtain insight into the density of enzymes around the substrate, we employed a template-matching procedure, as conducted above (Fig. 3B, orange). The enzymes were found in close proximity to the cellulose and appeared to penetrate between the cellulose chains to increase the efficiency of degradation. Indeed, our analysis revealed that the structure of cellulose undergoes major organizational changes during the interaction with the cellulosomes, transforming from a well-ordered cellulose structure (Fig. 3C) to deconstructed crystalline packaging (Fig. 3D). These novel observations provide snapshots of cellulose-degradation intermediates produced by the functional cellulosomal enzymes, located in the dense cellulosome layer around the bacterium, which further corroborated that our approach promoted the capture of real-time changes in cellulose organization and structure when interacting with cellulosomal complexes (Fig. 3C and D).

**Fig. 3.**
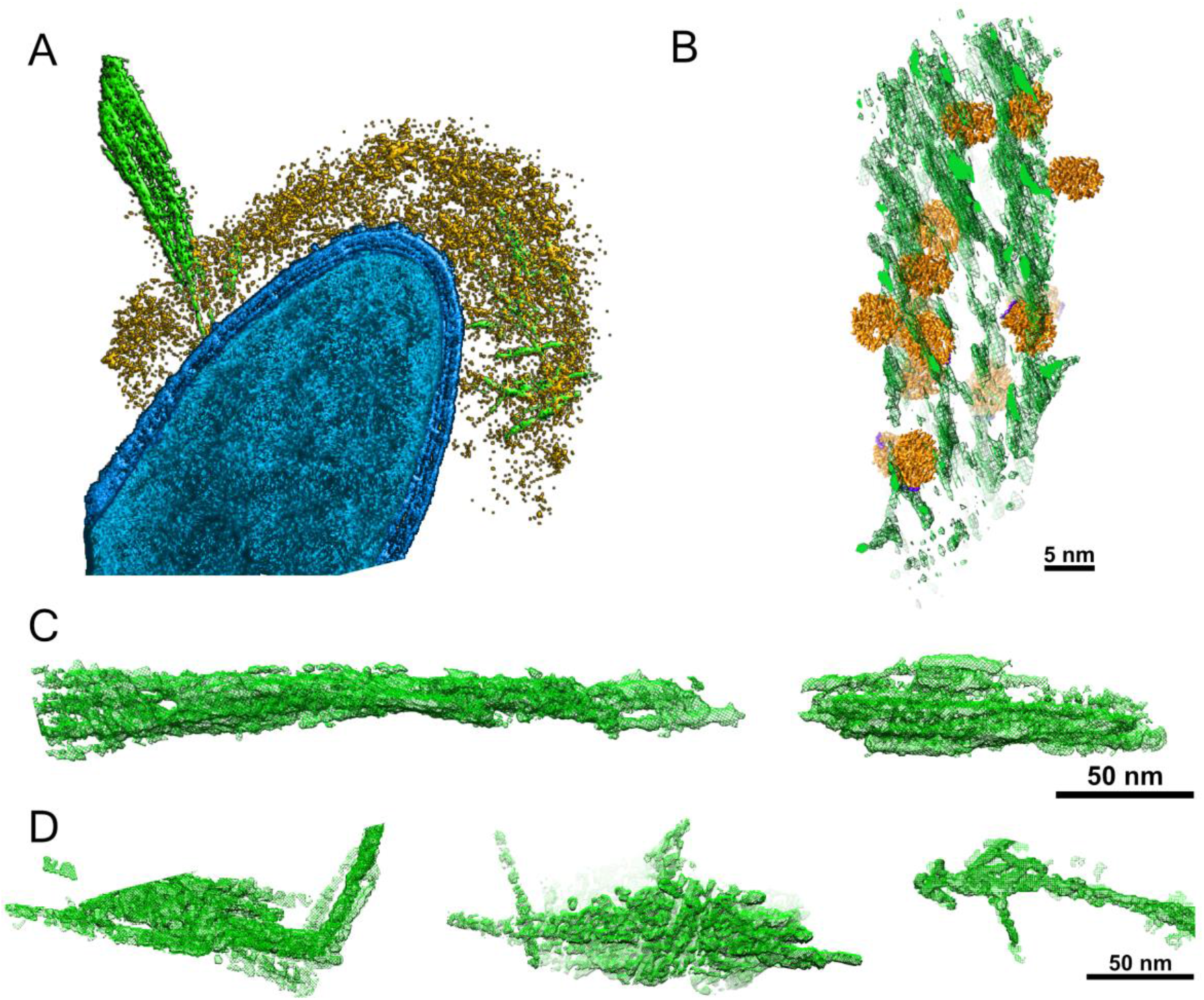
Interactions of cellulosomes with the cellulosic substrate. (A) Surface-rendered view of *C. thermocellum* (blue) shows the interactions of cellulosomes (yellow) with cellulose (green). Cellulose fibers are deconstructed into their component microfibrils by the enzymes within the cellulosome layer. (B) A rendered view indicating the position of cellulosomal enzymes around a cellulose fiber, enzymes (orange) and cellulose (green). Template matching of the Cel48S enzymes was applied to a subtomogram where cellulosome-cellulose interactions were observed. (C) Surface rendering of microcrystalline cellulose alone. (D) Surface rendering of cellulose fibers isolated *in silico* from positions in which interactions with cellulosomes were observed. The cellulosic crystal structures are altered (green) upon interaction with cellulosomes.

### Dynamic isogenic phenotypic heterogeneity of cellulosome organization

The detachment of the cellulosomal machinery during culture growth has been reported since its initial discovery ^4,41,42^, raising questions concerning the extent of this phenomenon and its single-cell distribution across the population as well as its ecological relevance. Indeed, while studying the structure of the cellulosomal machinery during cellulose degradation, we observed two distinct phenotypes of cellulosome organization on the bacterial cells, i.e., high-density and low-density organization (schematically shown in Fig. 4A). Interestingly we could not detect gradual decrease in density of the cellulosome layer around the bacteria in growing cultures (Supp. Fig. S6). To further pursue this phenomenon, we quantified the single-cell distribution of the cellulosome on the bacterial cells by employing cryo-EM as a tool for structural analysis of individual bacterial cells within the *C. thermocellum* bacterial population.

**Figure 4.**
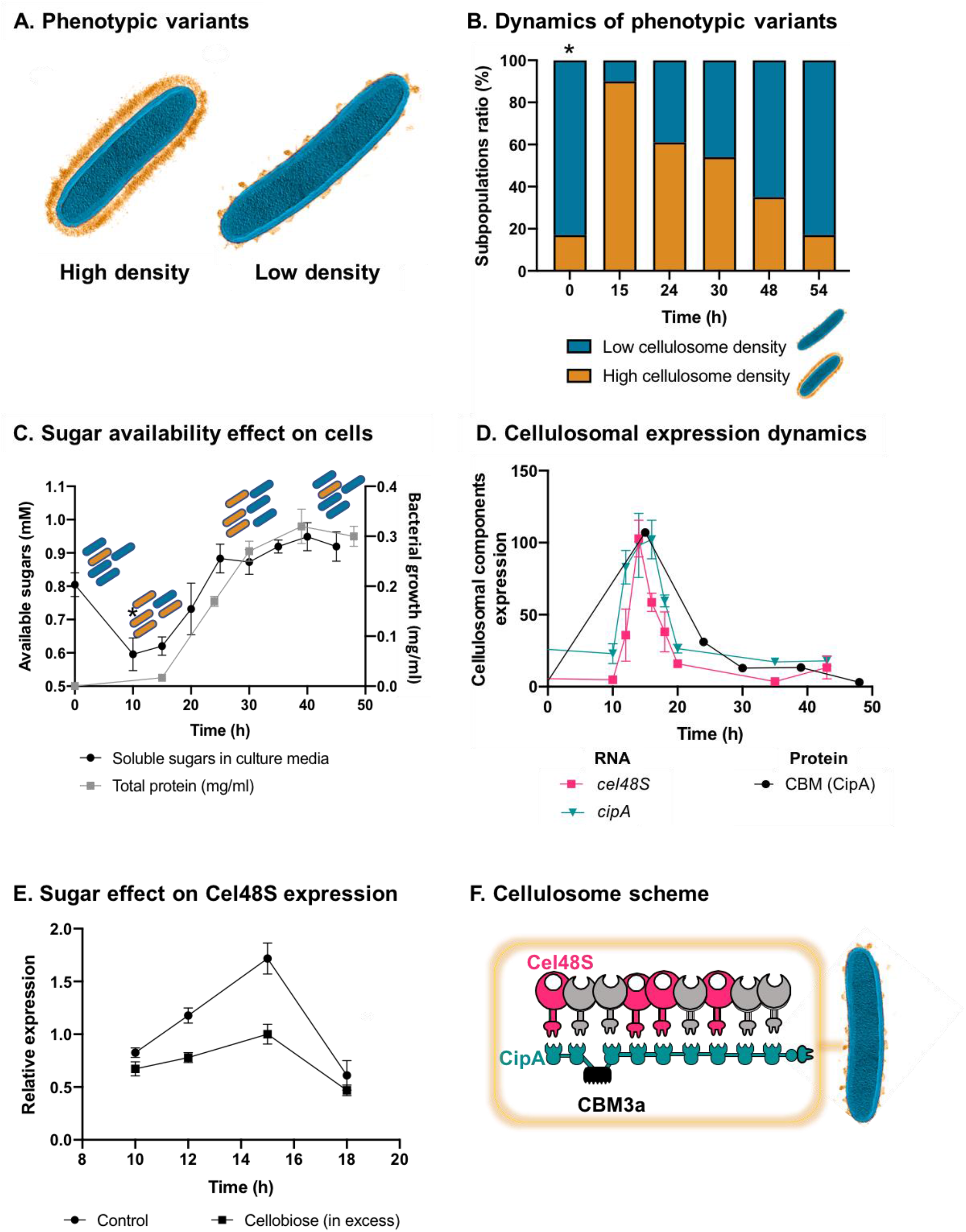
Dynamic isogenic phenotypic heterogeneity of cellulosome organization. (A) Schematic representation of cells from high- and low-cellulosome density subpopulations. (B) Dynamics of phenotypic variants in *C. thermocellum* subpopulations. A proportion of 5:1 (low-density:high-density) is observed during lag and stationary phases, while the high-density subpopulation is transiently dominant at 15 h, gradually decreasing in time to the initial proportion. Time 0 h was not measured, as it corresponds to the proportion of cells from the inoculum of time 54 h. (C) Available reducing sugars detected in *C. thermocellum* cultures grown on cellulose over time (black line, in mg/ml for total protein). A decrease in soluble sugars (10-15 h) is followed by a rise in their concentration, which is maintained stable until 50 h. Blue and orange rod-like shapes represent the low- and high-density subpopulations, respectively, and provide a schematic view of the relative changes in their distribution across the different growth phases and the levels of available sugars. (D) Expression dynamics of cellulosomal components at the level of transcript (RNA) and protein expression (units represent normalized copies/cells divided by 20,000 or band intensity, respectively). Per-cell RNA transcript copy number of *cel48S* (pink line) and *cipA* genes (blue line) from a culture of *C. thermocellum* growing on microcrystalline cellulose. At the protein level, CBM expression per cell was evaluated by using anti-CBM3a antibodies detecting the CBM present in the CipA scaffoldin. Cellulosomal gene expression is observed as a single peak after 15 h of growth of *C. thermocellum* in medium containing microcrystalline cellulose. (E) Cellulosomal gene expression is controlled by sugar availability. Addition of high concentration of cellobiose (10 g/L) into a 10-h culture of *C. thermocellum* negatively affects the expression of the main cellulosomal enzyme Cel48S (squares), compared to control (circles) as reported previously ^34^. (F) Schematic representation of the *C. thermocellum* cellulosome, highlighting the major components tracked in this study.

Our analysis revealed clear phenotypic heterogeneity within the same environment, under the same conditions with distinct cellulosome density distributions representing two different cell phenotypes (represented in Fig. 4A). The distribution of these two phenotypes in the stationary-phase culture shows a ratio of 1:5 (high versus low density). Phenotypic heterogeneity is considered to be an important ecological strategy for microorganisms to cope with environmental fluctuations and could be mediated by gene expression, where the proportion of cells that express the related phenotype changes as a function of environmental cues ^43^. To explore this notion, we subjected a stationary-phase culture, containing a large proportion of low-density phenotype, to a fresh medium, containing cellulose as the sole carbon source, and quantified the ratio between the populations of the two phenotypes during cell growth, using high electron-dose-exposure cryo-EM images (Fig. 4B). We analyzed 1160 individual cells, sampled from cultures at five time points after inoculation, which correspond to different population growth phases (with an average of n=232 cells per time point). Our analysis revealed that upon exposure of the bacterial cells to the new environment containing only cellulose, a sharp increase in the ratio of the high-density phenotype occurred, after which a gradual decrease in its concentration was observed back to a 1:5 ratio (Fig. 4B). This finding raised the hypothesis that soluble sugars, which are the product of the plant fiber degradation, are lacking at the beginning of culture growth, where only cellulose polymers are present as a carbon source. The soluble sugars accumulate during population growth and would affect the ratio between the two subpopulations. To explore this notion, we measured the availability of soluble sugars across the growth curve in the medium that contains cellulose as the sole carbon source. Our results revealed that the availability of soluble sugars derived from cellulose degradation is tightly connected to the ratio between the two subpopulations and culture growth stage (Fig. 4C). At early stages of population growth, where the availability of soluble sugars is low, an increase in the high-density subpopulation occurs, whereas, upon increase in sugar availability with population growth, the ratio of the low-density subpopulation increases (Fig. 4C). Interestingly, in the late stages of growth, when the high-density subpopulation comprises only 20-30% of the overall population, the amount of the soluble sugar concentration remained stable and did not decrease even when the population growth reached its peak. This observation supports the notion that soluble sugar is provided in saturating amounts at this stage, and the 2:10 ratio of the high-density sub-population is therefore maintained (Fig. 4C, 30 h to 50 h). These findings appear to reflect dynamic phenotypic variability, which infers that the accumulation of soluble sugars would be related to this phenomenon.

### Dynamic phenotypic variability is controlled by availability of soluble sugar

As our results suggest that the expression of the cellulosomal machinery changes during population growth and exposure to new environmental conditions, we proceeded to better understand cellulosomal expression dynamics and the factors that influence its regulation, within the context of our findings. To this end, we examined the expression dynamics and activity of the cellulosomal machinery during population growth after exposure to a fresh cellulose-containing media.

Our results showed similar dynamics to that of the single cell analysis observed by cryo-EM: i.e., a sharp increase of expression of the cellulosomal machinery upon exposure to the new environment, which was substantially reduced at the early stages of growth and maintained at basal levels. This increase in cellulosome expression was corroborated at the RNA levels of the two major cellulosomal components, Cel48S and CipA (Fig. 4D) corroborating previous reports ^44,45^, and at the protein level where we quantified the expression levels of CipA using a specific antibody against its CBM3a component (Fig. 4D). We also measured the total enzymatic activity normalized per cell of the culture, which comprises both the enzymatic machinery on the cell surface as well as secreted enzymes to the extracellular environment. The total cellulolytic activity presents the same pattern, which comprises a sharp increase in activity during early growth, followed by a sharp decrease in later growth stages (Supp. Fig. S7).

We next examined our hypothesis that soluble sugar availability resulting from cellulose degradation modulates the expression of cellulosomal components. To this end, we devised an experiment in which we added cellobiose, the major degradation product of cellulose, at the onset of cellulosomal expression (10 h after exposure of the cells to cellulose). Addition of cellobiose resulted in a negative effect on cellulosome expression (Fig. 4E), thereby suggesting that the availability of soluble sugars as products from the cellulose-degradation process is one of the cardinal ways for controlling the expression of cellulosomal components (Fig. 5).

**Figure 5.**
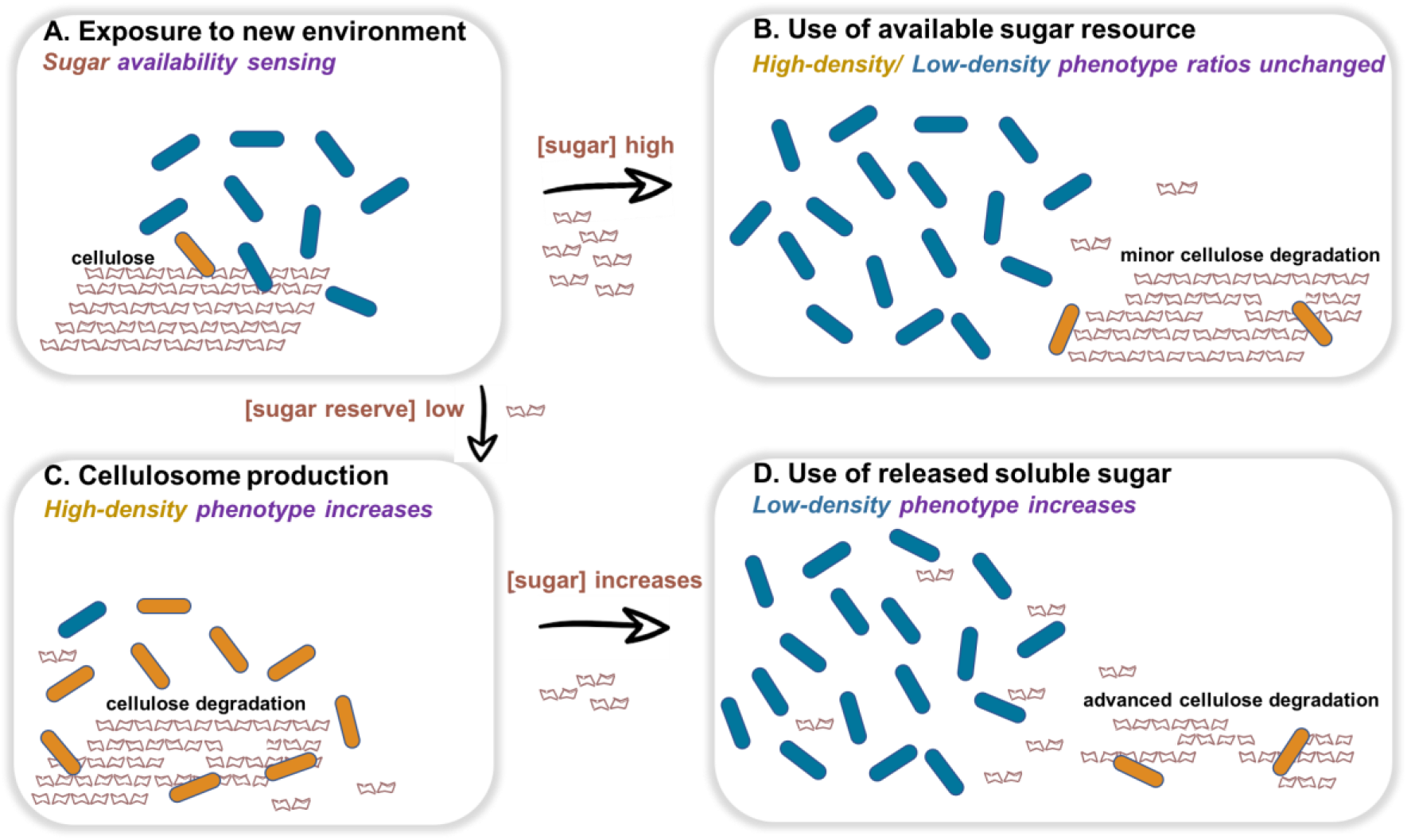
Dynamic phenotypic heterogeneity during cellulose degradation in the *C. thermocellum* population. The figure represents a schematic model of the findings revealed in our study. Blue and orange rod-like shapes represent the low- and high-density subpopulations, respectively. When *C. thermocellum* cells are exposed to a new environment containing cellulose (A), there are two possible outcomes depending on sugar availability. High sugar availability results in a basal level of cellulosome expression with no change in phenotypic heterogeneity of the population (B), where the high-density phenotype serves as a cellulosomal reservoir for future sugar-limiting conditions. In contrast, low sugar availability triggers the expression of the cellulosomal machinery (C), resulting in a major change in phenotypic heterogeneity of the population, where the high-density phenotype provides the present and future communities with high levels of sugar production from the cellulose substrate. When high sugar concentrations are available, the low-density phenotype can divide without producing the high-cost enzymatic machinery (D), and the high-density phenotype returns to low ratios of the population (bet-hedging strategy). The secreted cell-free cellulosomes and their interaction with the cellulose substrate are not represented.

## Discussion

In the present work, we studied the plant fiber-degradation process at the scale in which it occurs, i.e., micro to nanoscale level, from the individual microbes to the enzyme level, respectively. We selected *C. thermocellum*, one of the most efficient and characterized fiber-degrading microbes in nature. This bacterium deploys a cellulosomal strategy that allows it to extract energy from highly resistant plant-fiber polymers, notably crystalline cellulose.

Cellulosomes are extraordinarily efficient cellulosic fiber-degradation machineries that have evolved to provide sugar metabolites for subsequent assimilation by the parent bacterial cell. Our exploration spanned from unique mechanistic insights into the way cellulosomes are distributed on the bacterial cell wall, the way their enzymes interact with and degrade plant-fiber polymers, and the way single cells employ phenotypic heterogeneity at the population level.

Structural analysis by cryo-ET of *C. thermocellum* cells provided a tool to study the structure of cellulosomes in a near-physiological state, to understand their distribution on single cells, and to inspect the interactions of the bacterium with cellulose fibers (Fig. 3). In addition, our structure-based approach served to generate physiological and ecological hypotheses.

Our study revealed that cellulosome complexes are arranged as a highly dense interconnected layer of protein around the bacterial cell at a distance of ~60 nm from the cell wall. Intriguingly, we could directly observe the cellulosomal machineries during interaction with the plant-fiber polymers as they envelop them with their numerous enzymes and modulate the crystalline organization of the cellulosic substrate (Fig. 3).

We gained additional insights into the enzyme distribution on the cellulosome machineries by determining the structure of the dominant cellulosomal enzyme, namely the Cel48S cellobiohydrolase. Our data confirmed a previously published X-ray structure of this enzyme ^37^ and added new information regarding the amino acids that link the enzyme to its dockerin and on the interaction of this enzyme with cellulose. The structure of the Cel48S enzyme, determined up to 3.4 Å by cryo-EM, was used as an input for a template-matching approach to map the position of Cel48S around the bacterium. Our findings suggested that the exoglucanase spatially occupies the majority of the cellulosomal content under our experimental conditions, adding an additional dimension to previous proteomic studies, in which this protein was shown to be the most highly expressed enzyme of the cellulosome system ^33,35,36,39^. Moreover, this strategy allowed us to localize cellulolytic enzymes interacting with their substrate and revealed how they are arranged when attached to cellulose (Fig. 3B).

Our structural approach led to the detection and visualisation of two major subpopulations in the *C. thermocellum* culture under the same experimental conditions: one with cellulosomes covering the bacterial cells with very high density and another in which cellulosomes are nearly absent (Fig. 4A). We found that the ratio between these two phenotypes is dynamic and connected to the population growth stage and sugar availability (Fig. 4B and C). After exposure to new environmental conditions lacking soluble sugars, the expression level of the cellulosomal machinery increased in the majority of the cells in the population, after which it was repressed in most of the population. Furthermore, our findings indicate that the expression of the cellulosome machinery is triggered upon exposure to low levels of soluble degradation products (e.g., cellobiose) and then inhibited by increasing sugar availability (Fig. 4C and E). These findings add an additional dimension to the regulation of expression of the cellulosome machinery, which was reported previously to be mediated by sensing of extracellular polysaccharides via alternative sigma factors ^46^. In addition, it is interesting to note that the basal expression level of the cellulosome machinery at time 54 h is 20% of its highest expression levels at time 15 h (Fig. 4D). This corresponds to the proportion of the high-density subpopulation in the overall population (20%) at times 0 and 54 h (Fig. 4B), thus suggesting that the basal RNA levels stem from the expression of the cellulosome machinery in the high-density cell subpopulations.

The distribution of cellulosome density on the cell walls exhibited an all-or-none phenotype, where cells were either highly decorated with cellulosomes or almost devoid of cell-surface cellulosomes, without distribution of intermediate phenotypes (Supp. Fig. 6), thus suggesting that cellulosome shedding occurs in an abrupt manner similar to protein secretion and not a gradual process during growth. Taking our overall findings into account, it is tempting to speculate that a division-of-labor strategy takes place in the *C. thermocellum* population. In this scenario, the high-density subpopulation expresses the cellulosome machinery comprising enormous amounts of proteins and consequent metabolic costs, thus limiting the resources that this subpopulation can invest in growth, while at the same time the low-density subpopulation does not pay these costs and can invest them in propagation. Hence, the low-density subpopulation balances the essential but costly high-energy-process of expressing the cellulosomes by the high-density subpopulation.

Finally, our findings are consistent with a bet-hedging strategy in the *C. thermocellum* population, mediated by phenotypic heterogeneity, whereby the presence of diverse phenotypes adapted to different environments would spread the risk of varying environmental conditions ^47^. Such a strategy would be effective for organisms under fluctuating environmental conditions, where a proportion of individuals in the population pay an energetic/fitness cost by “preparing” for changing environmental conditions. Similarly, in our case, although the soluble sugar levels are not limiting (Fig. 4C), the high-density cell subpopulation, albeit in lower ratios, still expresses the costly cellulosomal machinery and would therefore provide the first line of defense for a future nutrient-poor and hostile environment containing only cellulose polymers. Our approach, whereby structural biology under near-physiological conditions was employed to understand microbial ecology, provides a pioneering view, at the micro and nano scales, into the unique organization of the cellulosome machinery on the bacterial cell surface, the way cellulosomes interact with their cellulosic substrate and how they are distributed and regulated across the bacterial population during the growth process.

## Authors’ contribution

M.T, S.M, O.M and I.M conceived and designed the experiments; M.T acquired and analysed the cryo-EM and cryo-ET data, S.M and O.T performed the microbial and biochemical work. O.M and IM secured funding for the project. M.T, SM, IM and OM wrote the manuscript with contributions from all of the authors.

## Data availability

Structural data that support the findings of this study on the structure of Cel48S have been deposited in the Electron Microscopy Data Bank under accession code EMD-11986. Cryo-ET data have been deposited EMPIAR-10593.

## Acknowledgements

This work was funded by the DIP (2476/2 −1) to I.M and O.M., ERC (64084, to I.M), Swiss national foundation grant (SNSF 31003A_179418) to O.M. SM is grateful to EMBO for the short-term fellowship (7686) allowing a visit to the University of Zurich. The authors thank Eva Setter-Lamed (Weizmann Institute of Sciences, Israel) for technical assistance, Matthias Wojtynek (UZH/ETH) for assisting with template matching procedures, Rebecca de Leeuw (UZH) and Miriam Weber (UZH). The authors thank the Center of microscopy and image analysis (ZMB) at the university of Zurich, Ilse Katz Institute for Nanoscale Science and Technology Shared Resource Facility under the direction of Dr. Uzi Hadad for image acquisition with a Zeiss LSM880 Airyscan.

## Materials and Methods

### Expression of the Cel48S enzyme

The plasmid for Cel48S expression was previously described ^48^. Competent *E. coli* BL21 Star (DE3) cells were used to express the protein in 2-L cultures, using isopropyl-1-thio-β-D-galactoside (IPTG) (Fermentas UAB Vilnius, Lithuania) to induce expression. Cells were harvested by centrifugation at 3000 *g* for 5 min, and the protein was purified using a Ni–NTA column (Qiagen, Hilden, Germany) as reported previously ^49^. The protein was further purified by size exclusion chromatography (Superdex 200 Increase, GE Healthcare Biosciences, Piscataway, NJ) in TBS (137 mM NaCl, 2.7 mM KCl, 25 mM Tris at pH 7.4) +2.5 mM CaCl_2_ + 2 mM *β*-mercaptoethanol buffer.. The expected peak was eluted at 12 ml, and the molecular weight of the protein in the selected fraction was confirmed by SDS-PAGE and mass spectrometry.

### Sample preparation for cryo-EM and tomography

The concentration of Cel48S was diluted to ~0.35 mg/ml in ice-cold elution buffer, before being vitrified on carbon-coated Au grids (R06/1, 200 mesh; Quantifoil, Jena, Germany).

*C. thermocellum* DSM1313 cells were grown on GS2 medium^36^ containing either 8% cellobiose or 0.2% microcrystalline cellulose as carbon source. The cells were grown anaerobically at 60°C. The cells grown on cellobiose were harvested at 24 h, and the cells grown on crystalline cellulose were harvested at different time points, ranging from 15 h to 54 h of incubation. Cells were centrifuged for 5 min at 5000 *g*, quickly washed in TBS buffer, and resuspended in TBS prior to flash-freezing. A volume of 3 μl of cell sample was mixed with 3 μl of 10 nm fiducial BSA gold marker (Aurion, Wageningen, The Netherlands) and applied on glow-discharged copper EM grids, coated with a carbon mesh (R2/1, 200 mesh; Quantifoil, Jena, Germany). The grids were manually blotted for ~4-5 s from the reverse side and plunge frozen in liquid ethane by using an in-house plunger.

To analyse the organization of microcrystalline cellulose (Sigma-Aldrich), dilute suspensions (0.04-0.0016 g/l in TBS) were prepared, 3 μl of the suspensions were mixed with 3 μl of 10 nm fiducial BSA gold marker (Aurion, Wageningen, The Netherlands) and applied on glow-discharged copper EM grids, coated with a carbon mesh (R2/1, 200 mesh; Quantifoil, Jena, Germany).

### Cryo-EM data acquisition

A total of ~2000 micrographs were acquired using Titan Krios (Thermo Fisher Scientific, Waltham, MA) equipped with Gatan energy filter and K2 summit (Gatan Inc., Pleasanton, CA). Using an underfocus range of 0.5-1.5 μm and the super resolution mode resulted in a pixel size of 0.425 Å. Exposures of 0.2 *s* for 12 *s* resulted with 60 frames and a total dose of ~67 e^-^ /Å^2^.

### Cryo-EM image processing

The micrographs were drift-corrected and binned once using Relion 3.0, resulting in a pixel size of 0.86 Å. A total of ~1100 manually selected particles were used to create a 2D reference for further automatic particle picking using Relion 3.0. The structural determination steps are described in Figure S4 and Table 1. Initially, a few rounds of reference-free 2D classifications using ~ 363 000 particles resulted in an initial reference using Relion 3.0 by Stochastic Gradient Descent. 3D classifications, and 3D refinements were performed by using the initial reference with the refined 3D classes. One the 3D-refined class reached 3.6-Å resolution and was further improved by Bayesian polishing and CTF refinements. The final 3D structure was determined to be 3.38 Å resolution, using ~ 55 000 particles.

The obtained final model was compared to the pdb of the Cel48S exoglucanase (5yj6) by the fit-in-map option in Chimera, resulting in a correlation score of 0.87. The stoichiometry of the model was improved by real-space refinement in Phenix, and the final atomic structure was validated by MolProbity.

### Immunogold labelling of cellulosomes

The protocol was adapted from ^50^. A 1-ml sample of bacterial culture was washed in TBS as described above, and 5 μl of the washed cells were introduced to Au grids (R2/1, 200 mesh; Quantifoil). The grids were fixed in 4% PFA (in TBS) for 5 min at room temperature and were quenched in 0.05 M glycine/PBS for 15 min at room temperature to inactivate aldehyde groups present after aldehyde fixation. Next, samples were blocked in blocking solution (5% bovine serum albumin (BSA)/0.1% cold water fish skin gelatin/PBS; Aurion) for 30 min at room temperature, and subsequently washed with the incubation solution (0.2% BSA-c/PBS; Aurion). A volume of 20 μl of anti-CBM antibody ^51^ (1:1000 dilution in incubation solution) was incubated on grids for 1 h at room temperature. The EM grids were washed with the incubation solution before treatment with gold conjugate (Protein A-6 nm gold; Aurion) Finally, the grids were washed extensively in PBS (3×5 min),10 nm fiducial gold marker solution was added prior to vitrification in liquid ethane cooled liquid N2.

### Cryo-ET data acquisition and reconstruction

Sixty tilt-series of 24-h grown *C. thermocellum* bacteria cultured on microcrystalline cellulose were acquired at 53,000 magnification corresponding to a 0.27 nm pixel size, at a defocus of −1 μm. The tilt series are covering an angular range of −60° to +60° with an angular increment of 2° and total dose ~150 e-/Å^2^, using a Thermo Fisher Titan Krios electron microscope operating at 300 keV, equipped with a quantum energy filter, a K2-Summit direct electron detector (Gatan Inc.) and a Volta phase plate (VPP, Thermo Fisher Scientific, Eindhoven, Neaderlands).

Tilt series of Microcrystalline cellulose were acquired at 53 000x magnification, covering an angular range of −60° to +60° with an angular increment of 2° and total dose ~150 e-/Å^2^ and at −6 defocus for visualization purposes.

Interactions of microcrystalline cellulose with cellulosomes were analysed using *C. thermocellum* cultures grown for 15 h; 23 tilt-series at 53 000x magnification and a defocus of 1-2 μm were acquired using VPP. A single tilt-series was recorded using an under-defocus value of 6 μm for visualization purposes (Fig. 3A). Additional 25 tilt-series of 15-h grown *C. thermocellum* bacteria cultured on cellulose were acquired at 105,000x magnification, corresponding to a 0.13 nm pixel size, at an under focus of 4 μm, and an angular increment of 3°. All tilt-series were reconstructed by using the IMOD workflow. Subvolumes were produced by the post-processing trimming option of IMOD.

For population analysis, 4800x-magnification images of *C. thermocellum* incubated for five different time periods 15 h (42 images), 24 h (219 images), 30 h (69 images), 48 h (40 images) and 54 h (231 images), were acquired.

### Image Processing

#### Population analysis

The total number of high-cellulosome and low-cellulosome expressing bacteria (non-lysed and non-sporulating) were manually counted in low-magnification images of *C. thermocellum* for different incubation periods, 84 bacteria for 15h, 499 bacteria for 24h, 179 bacteria for 30h, 170 bacteria for 48 h, and 230 intact bacteria for 54 h. The total number of high-cellulosome density maintaining bacteria was also counted manually for the same images, and percentages of high-cellulosome-density-maintaining bacteria were calculated for each condition in an Excel document. The final histogram of the percentages for each condition was plotted in MATLAB (Fig.3C).

#### Thickness measurements

Central x-y slices, ~70 nm in thickness, through 15 bacteria were projected onto a plane. The images were exported to the Fiji software package. The thickness of the given cellulosome layer was measured by subtracting the length from the bacterial S-layer to the proximal cellulosome layer edge from the distal edge of the cellulosome layer. The measured lines were selected to be perpendicular to the S-layer plane. The measurements from each tomogram were analysed by Excel, Fig. S3B. The distance from S-layer to the proximal edge of the cellulosome layer was measured accordingly (Fig. S3C).

### Template matching

*In situ* Cel48S enzymes were localized by PyTom template matching in tomograms and a subvolume ^52^. *In vitro* Cel48S structure was filtered to 2.2 nm, applied as previously ^53^. The assigned particles were inspected manually in Amira, the coordinates on the bacteria and false positives were removed by creating a mask that excludes them. Later, the coordinates inside the masked region were removed by an in-house script developed in MATLAB. The final number of assigned Cel48S units after masking were shown in Fig.S7 for each of 8 high-density bacteria tomograms, respectively, and 63 units for the subvolume (Fig.3D).

### Mapping the Cel48S structure into tomographic volumes

Randomly oriented *in vitro* Cel48S structure was used to map back into the tomograms of the bacteria. The distances between the center of mass of neighbouring structures-coordinates-were calculated and plotted in MATLAB (Fig2 F).

For visualizing enzyme densities attached to cellulose (Fig. 3B), the assigned angles obtained from the template matching procedure were used displaying the *in vitro* Cel48S structure, at the cellulose surface. Figure 5B was made using Chimera software (UCSF, USA).

### Segmentation

The cellulosomes and bacteria were segmented by the isosurface thresholding option in Amira-Avizo 2019.1 (Thermo Fischer Scientific).

### Confocal microscopy

A volume of 2 ml of cells grown with microcrystalline cellulose was centrifuged for 5 min at 5000 *g*. The cells were washed quickly in TBS and fixed with paraformaldehyde (Sigma-Aldrich, St. Louis, MO) diluted to 4% in TBS for 20 min. The cells were then washed and centrifuged for 5 min at 5000 *g*, three times with TBS and once in TBS containing 1% Triton X-100 (Sigma-Aldrich). The cells were then re-suspended in 5% BSA diluted in TBS for blocking 1 h at room temperature. After removing the blocking solution by centrifugation, rabbit anti-CBM3a from CipA antibody diluted at 1/5000 in 1% BSA in TBS was added, and the cells were incubated 1 h at room temperature. The cells were washed three times with TBS by centrifugation as described above, and secondary goat anti-rabbit IgG Alexa 594 antibody (Thermo Fisher Scientific) diluted at 1/1000 in 1% BSA in TBS was allowed to interact for 1 h at room temperature. After three washes of the cells (by centrifugation as above), the cells were mounted on slides with SlowFade^®^ Gold anti-fade reagent (Thermo Fisher Scientific) using polylysine-coated cover slides. Confocal images were obtained by using a LSM880 inverted laser-scanning confocal microscope (Carl Zeiss AG, Oberkochen, Germany) equipped with an Airyscan high-resolution detection unit. A Plain-Apochromat 63x/1.4 Oil DIC M27 objective with a 561 nm DPSS laser with a no-emission filter was used, and parameters were set to avoid pixel intensity saturation and to ensure Nyquist sampling in the XY plane.

### RNA analysis

Three independent sets of cellulose-grown cell cultures were individually sampled at different growth times from 10 to 43 h. Cells were harvested for 5 min at 3000 *g*, and the pellets were resuspended in *RNAlater* solution according to the manufacturer’s protocol. Cells were then stored in −80°C until downstream RNA extraction preparations. The RNA was isolated using an RNAeasy minikit (Qiagen, Germantown, MD) using protocol 4 with the following modifications: cells were incubated 2 h at room temperature in TBS prepared with RNase-free water supplemented with lysozyme at a concentration of 15 mg/ml and 20 μl proteinase K. Then the purified RNA samples were immediately processed with the Thermo Fisher Scientific verso cDNA synthesis kit.

Quantitative real-time PCR analysis was performed to calculate the RNA levels of two genes of interest: Cel48S and CipA RNA, which were normalized to RecA RNA levels. A specific fragment of each gene was amplified using the primer pairs: 5’-CGCAGAAGGCCGTGCTATA-3’ and 5’-CAGAACCTTTACCCTGCTCCTTT-3’ for Cel48S, 5’-CAGTATGCTCTTAGTTGTGGCTATGC-3’ and 5’-TGATCCAACGGCTGCTGTAA-3’ for CipA, 5’-GTTGCGGTAAATCTCGATATTGTAAA-3’ and 5’-GGCCAATCTTCTGACCGTTG-3’ for RecA. For each gene, individual standard curves were generated by amplifying serial 10-fold dilutions of quantified gel-extracted PCR products obtained by the amplification of each fragment. The standard curves with required standard efficiencies were obtained using six dilution points and were calculated using the Rotorgene 6000 series software (Qiagen). Realtime PCR was performed in a 10 μl reaction mixture containing 5 μl Absolute blue SYBR green master mix (Thermo Fisher Scientific), 0.2 μl of each primer (diluted to 10 μM), 3.6 μl nuclease-free water, and 1 μl of cDNA.

### CipA protein expression analysis

A volume of 2 ml samples from the three independent sets of cellulose-grown cell cultures was taken from times 15 h, 24 h, 30 h, 39 h, 48 h and 54 h. Half of the volume was used to determine the cell numbers in the culture, by DNA extraction as performed by Stevenson and Weimer ^54^ without the initial filtration steps, followed by quantitative real-time PCR analysis as described above using 16S universal primers covering the V2 and V3 regions of the gene (5’-TTTGATCNTGGCTCAG-3’ and 5’-GTNTTACNGCGGCKGCTG-3’).

The calculated number of 16S copies served to normalize the volume taken for Western blotting to a similar number of cells. Normalized volumes of samples from the different growth time points were applied on SDS-PAGE gels (10% acrylamide) and transferred to a nitrocellulose membrane using Mini Trans-Blot cells (Bio-Rad Laboratories Ltd., Rishon Le Zion, Israel). Nonspecific protein interactions were blocked by incubating the membrane for 1 h with 5% BSA diluted in TBST (Tris-buffered saline with 0.05% of Tween 20). Rabbit antibody against CBM3a from CipA ^51^ at a dilution of 1:3000 was incubated with the nitrocellulose membrane for 1 h in 1% BSA diluted in TBST. The membrane was then rinsed three times with TBST and then incubated for 1 h with secondary anti-rabbit antibody labeled with horseradish peroxidase (HRP) at a dilution of 1:10,000. The membrane was rinsed again three times with TBST and developed by incubation for 1 min with equal amounts of solutions A and B (ECL, Ornat Biochemicals & Lab Equipment, Rehovot, Israel). Chemiluminescence was quantified using a luminescent image analyzer (Fusion, FX7 G/R/IR).

### Cellulolytic activity per cell

An overnight culture from *C. thermocellum* was used as a starter culture to inoculate 125 mL serum bottles containing 100 mL of GS2media with microcrystalline cellulose as the sole carbon source. After inoculation, the serum bottles were incubated at 60°C for 45 h and samples were taken at 0, 10, 15, 20, 25, 30, 35, 40 and 45 h. The samples were centrifuged at 4000 rpm for 5 min to separate the bacterial pellet from the supernatant, and the protein fraction was precipitated by adding ammonium sulphate at 80% saturation in order to recover the cellulosomes that were released into the media. Proteic pellets were obtained by centrifuging the solutions from ammonium sulphate treatment at 4000 rpm for 10 min, resuspending the pellets in 1 mL of TBS buffer and then reuniting the resultant solution with the bacterial pellet collected from the original sample (overall x15-fold concentration). Then, these samples were used for determining the cellulolytic activity per cell using phosphoric acid swollen cellulose (PASC) as a substrate ^55^. Enzymatic reactions were performed at 60°C for 72 h under continuous agitation (600 rpm). After incubation, samples were centrifuged at 14,000 rpm for 10 min, and the supernatant was used to determine the amount of released sugars, by the DNS method ^56^. Cell numbers in each sample from different time points were normalized by absolute quantification of the 16S gene copy number using real-time qPCR.

### Effect of soluble sugars on CipA and Cel48S gene expression

In order to determine the effect of soluble sugars on the expression of CipA and Cel48S genes in *C.thermocellum*, either cellobiose or sugars released from microcrystalline cellulose by a 20-h *C. thermocellum* culture were added into the culture media. In the case of cellobiose, a 10-h growth culture of *C. thermocellum* was supplemented with 10 mL of cellobiose 10% w/v and samples were taken after 10, 12, 15 and 18 h. Alternatively, taking into account that the peak of expression of the cellulosomes occurs at ~15 h after inoculation, we extracted samples of 0.45-μm filter-sterilized supernatant from a 20-h growth culture and used it as the culture media to inoculate the total cells of a 10-h culture, obtained by centrifuging the cells (and cellulose) for 10 min at 4000 rpm. Before inoculation, cells were washed twice in fresh medium, in order to remove soluble sugar. All the steps were performed under anaerobic conditions. The cultures were incubated at 60°C, and 2 mL samples were taken at 10, 12, 15 and 18 h. Finally, RNA was analysed as described above.

## Notes

### Competing Interest Statement

The authors have declared no competing interest.

