## Supplemental figures&table for "Nanoscale resolution of microbial fiber degradation in action": Nanoscale resolution of microbial fiber degradation in action_supplementary.pdf

**Supplementary figures**

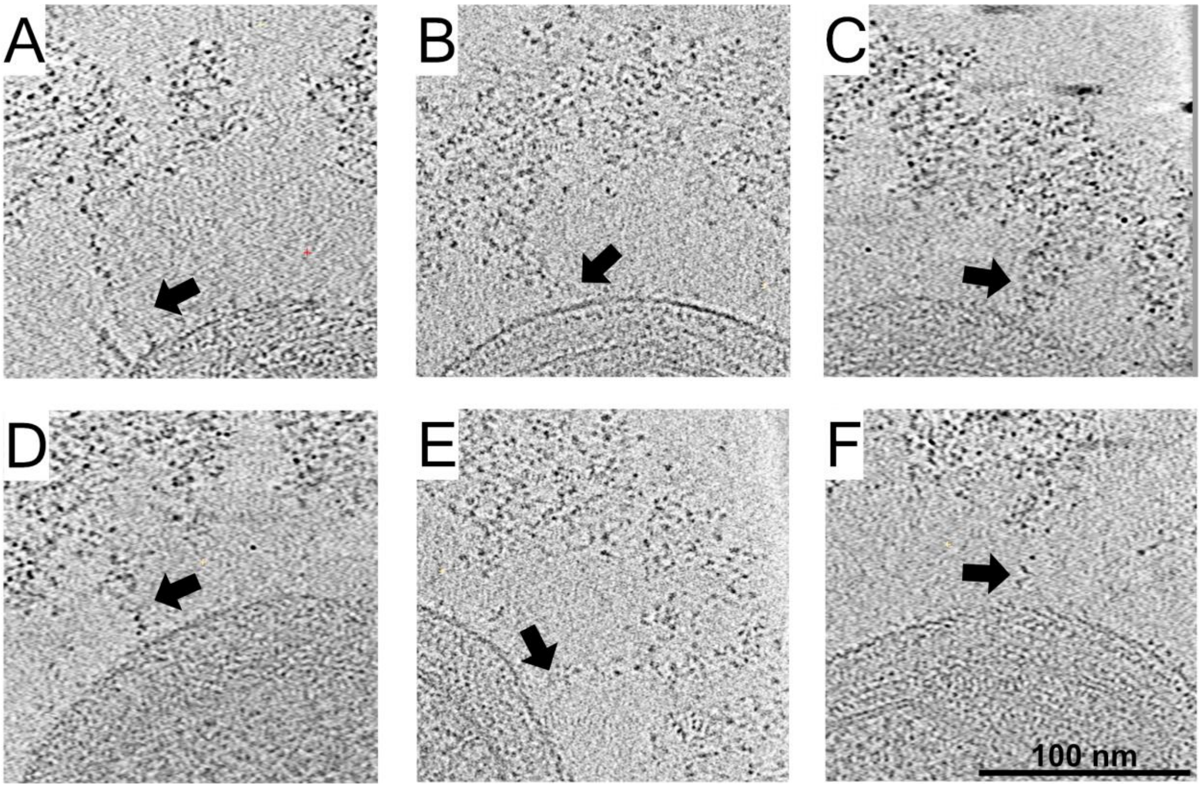


**Fig.S1. Cellulosomes emanate from the cell wall towards the densely packed layer.** A density span between the bacterium and the cellulosome layer (arrows) was detected. x-y slices through six bacteria, representing 5.4 nm in thickness.

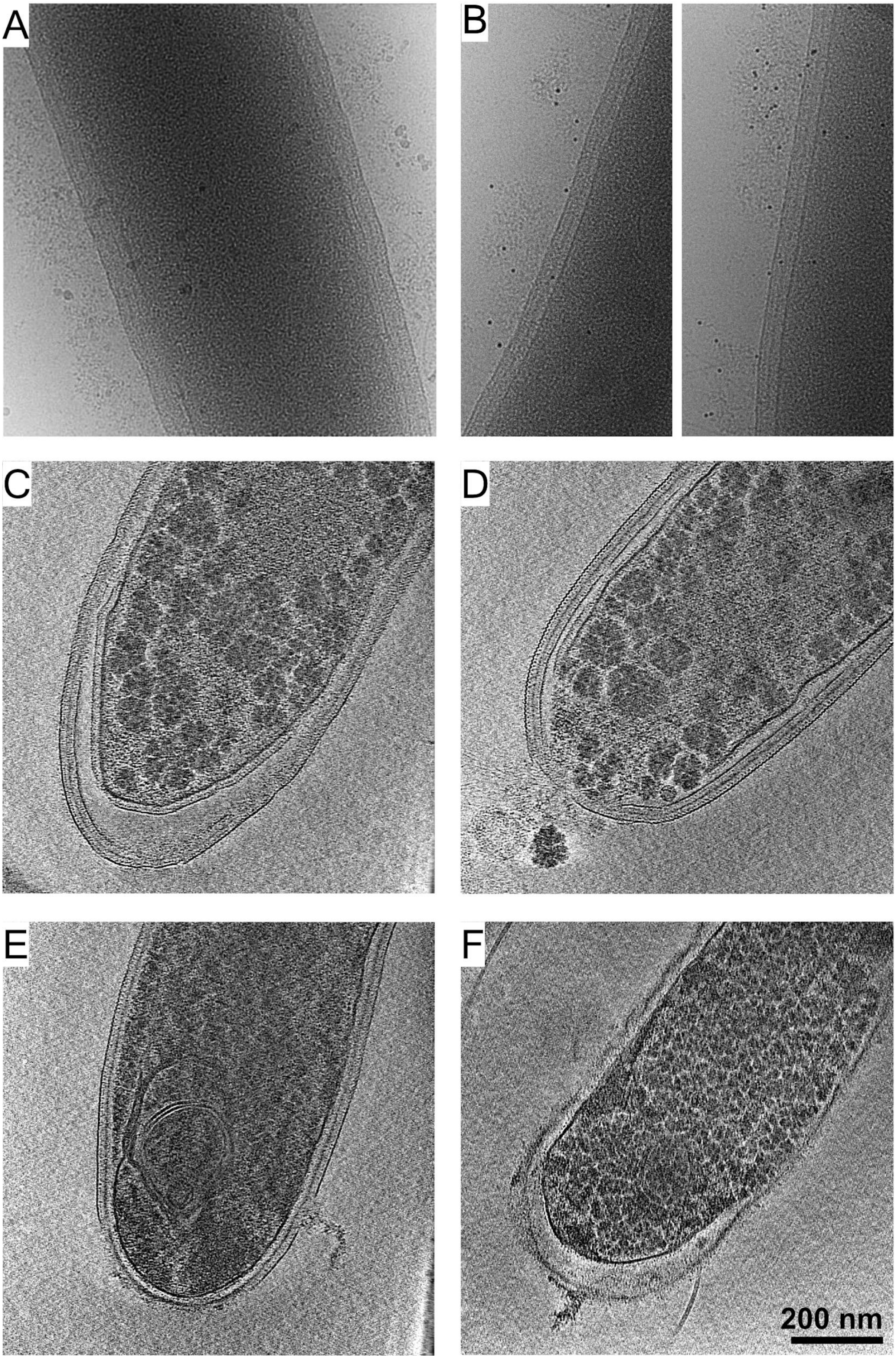
A
B
C
D
E
F
200 nm

**Fig. S2. The cellulosomal protein layers around *C. thermocellum* cells.** Immunogold labeling using protein A coupled with coloidal-gold showed no observed signals (A), while the addition of the anti-CBM3a antibody directed the localization of the gold-conjugated protein A to the proteinous layer around the bacterium (B). Cryo-ET analysis of mutant strains: (C and D) Strain CTN7 expresses none of the major scaffoldins (i.e., CipA, OlpB, Orf2p, SdbA, OlpA and OlpC). (D and F) Strain DS11 lacks the major enzyme-integrating scaffoldin (CipA). Neither strain produces cellulosomal structures outside the cells. These experiments indicate that the observed exogenous layer corresponds to cellulosome related proteins.

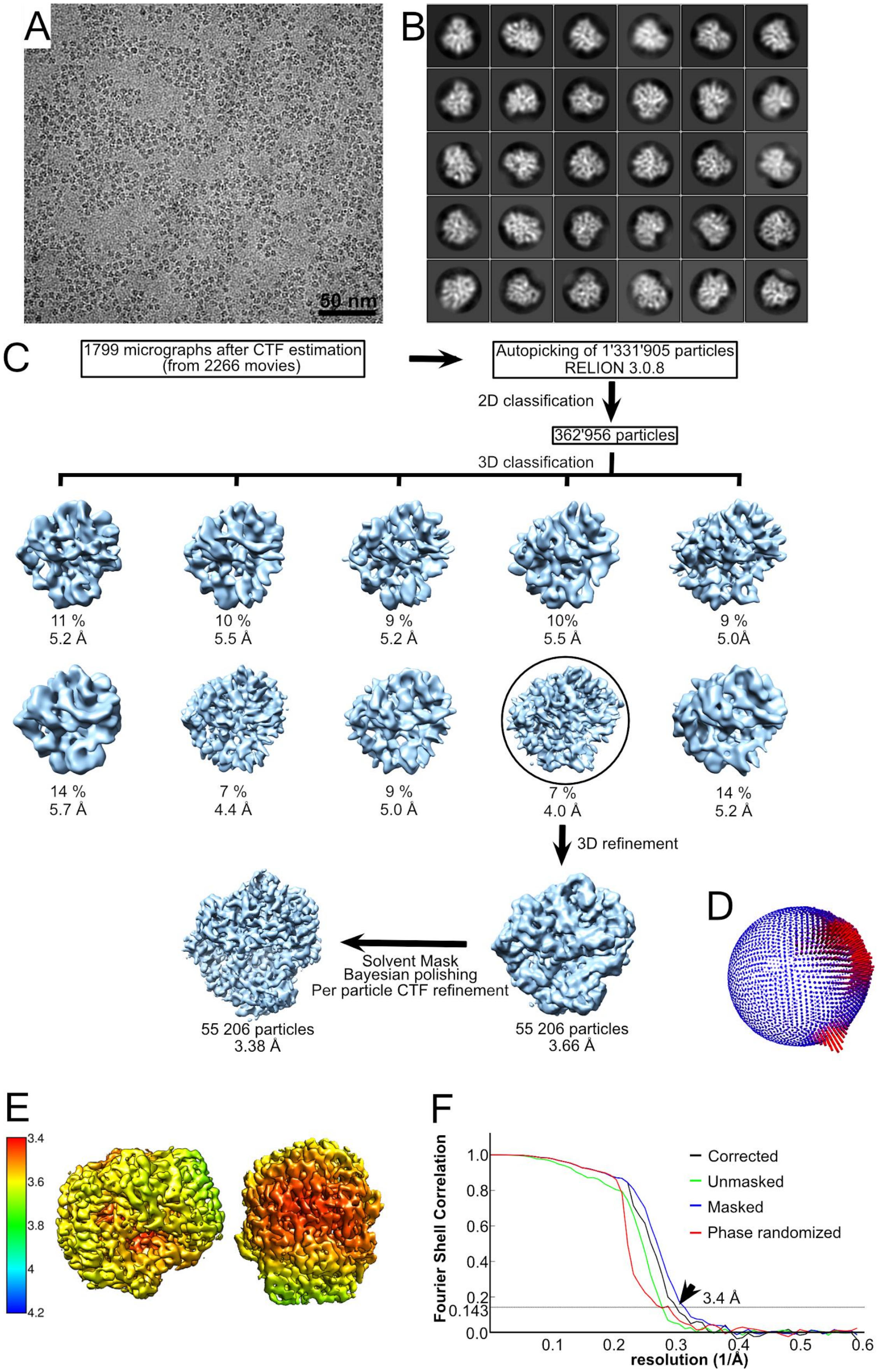
A
50 nm
B
C
1799 micrographs after CTF estimation
(from 2266 movies)
Autopicking of 1'331'905 particles
RELION 3.0.8
2D classification
362'956 particles
3D classification
11 %
5.2 Å
10 %
5.5 Å
9 %
5.2 Å
10%
5.5 Å
9 %
5.0Å
14 %
5.7 Å
7 %
4.4 Å
9 %
5.0 Å
7 %
4.0 Å
14 %
5.2 Å
3D refinement
Solvent Mask
Bayesian polishing
Per particle CTF refinement
55 206 particles
3.38 Å
55 206 particles
3.66 Å
D
E
3.4
3.6
3.8
4
4.2
F
Fourier Shell Correlation
resolution (1/Å)
Corrected
Unmasked
Masked
Phase randomized
3.4 Å
0.143
0.0
0.2
0.4
0.6
0.8
1.0
0.1
0.2
0.3
0.4
0.5
0.6

**Fig. S3. Structural determination of the Cel48S enzyme.** (A) A typical micrograph depicting the major cellulosomal exoglucanase, Cel48S. (B) The final 2D classification showing different orientations of this asymmetric protein. (C) The workflow of image processing for Cel48S. After several rounds of 2D classifications removing false positives, 362,956 particles were selected for initial 3D model building. The particles were subjected to a couple rounds of 3D classifications and refinement. The percentage of particles within the resultant 3D classes and the respective resolution of the 3D classes are visualized. Further refinement with 55,206 particles is shown in the black circle. After final refinement with B-factor of −110 Å$^2$, a map of 3.66 Å was resolved. The map was post-processed with a solvent mask, and the particles were subjected to Bayesian polishing and CTF refinement. These operations improved the map to 3.38 Å. (D) The angular distribution of all 55,206 particles that contributed to the final Cel48S map is depicted. The height of the cylinder bars (from blue to red) are correlated with the number of particles in this view. (E) Local resolution variations in the cryo-EM map. The resolution ranges from 3.4-3.8 Å as calculated by the Relion local resolution option. (F) Fourier-shell correlation (FSC) plots of the final Cel48S density map. The plot shows the unmasked (green), masked (blue), phase randomized (red) and masking-effect-corrected (black) FSC curves. The resolution at which the gold-standard FSC curve drops below the 0.143 threshold is indicated (arrow).

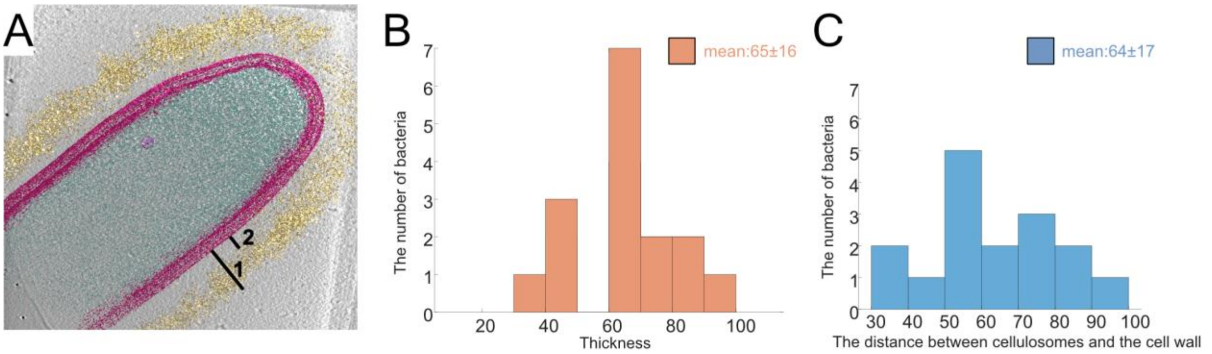


**Fig. S4. Position and thickness of the cellulosome layer.** The thickness of the cellulosome layer (A:1-2) and the distance between the cell wall and cellulosomes (A:2) were measured around the bacteria (B,C, respectively, see Materials and Methods Section). Analysis of 15 tomograms was used.

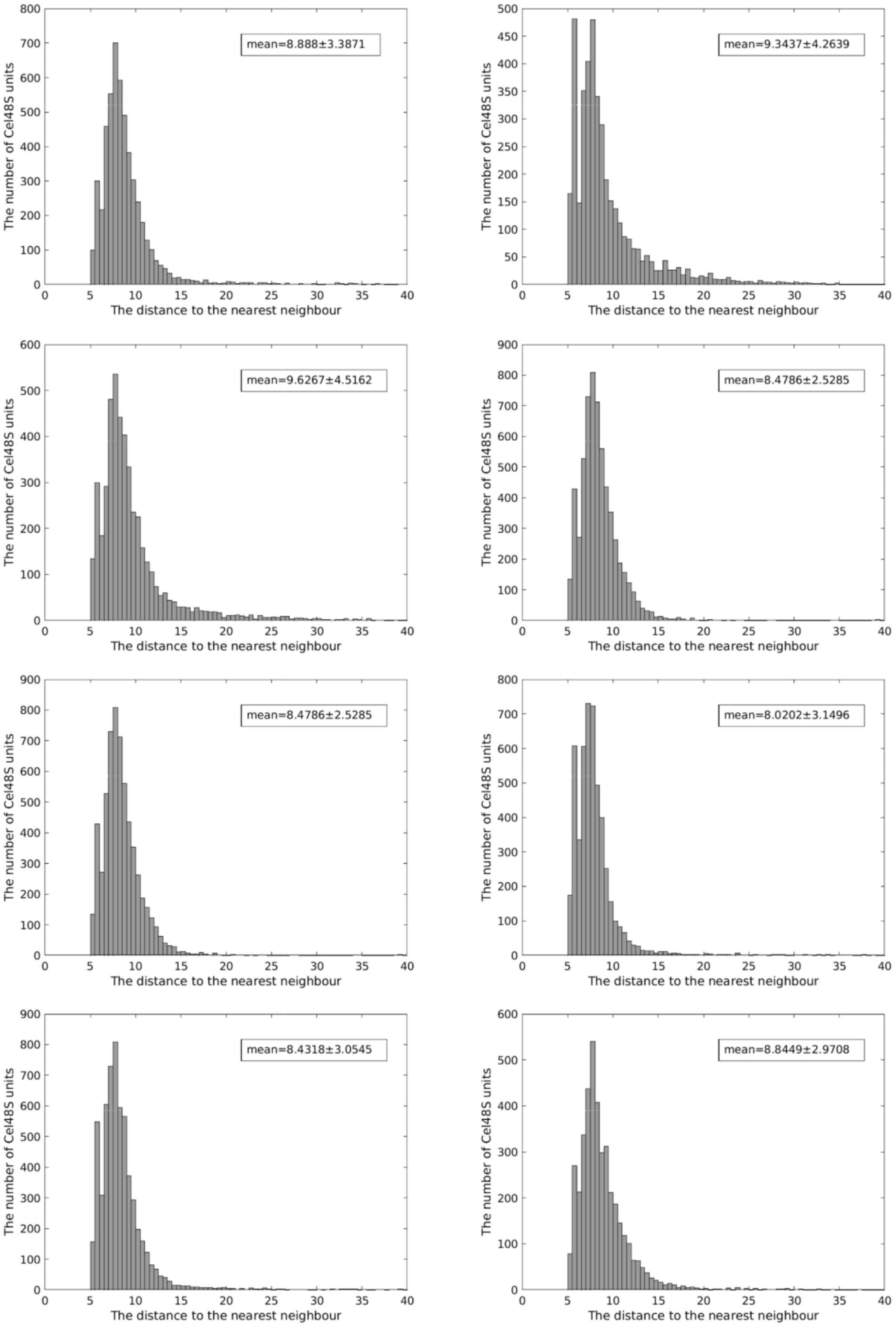


**Fig. S5.** Nearest-neighbour analysis of detected Cel48S units in 8 tomograms of high-density bacteria. The mean distance to the nearest neighbour changes between 8 and 9.6 nm.

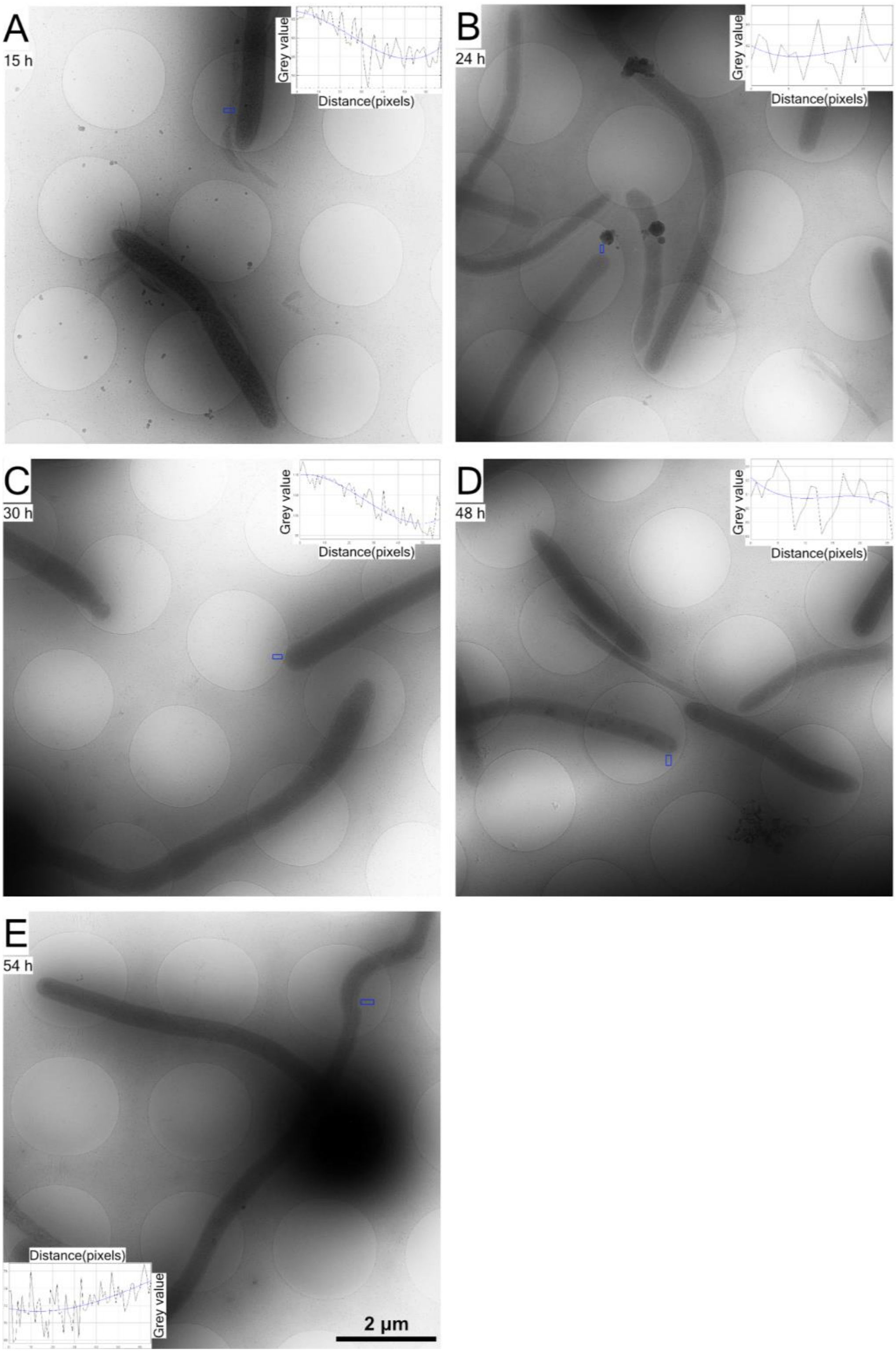


**Fig. S6. Representative cryo-EM images of bacterial cultures.** Sampling bacterial cultures at specific time points (A-E, 15, 24, 30, 48 and 54 h, respectively). Examples of intensity profiles of the regions within blue rectangles near bacteria were plotted. The lower values in the plots represent the denser regions of the image. In the presence of the dense cellulosome layer, sharp decrease in grey values were observed in the plots.

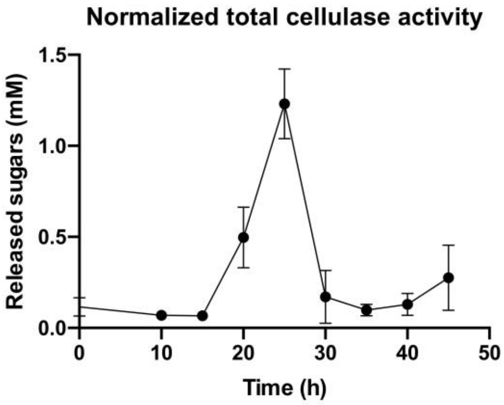


**Fig. S7. Overall enzymatic activity of the cellulosomal machinery during population growth**. A sharp increase in the total cellulase activity normalized by cell number is observed at the early stages of bacterial growth and is followed by a marked decrease at the end of the exponential phase.

**Table 1: Data collection and processing statistics of the cryo-EM structure**

| | |
|---|---|
| **Data collection** | |
| Microscope | FEI Titan Krios |
| Voltage (kV) | 300 |
| Camera | Gatan K2 Summit |
| Camera mode | Super-resolution |
| Energy Filter | Gatan postcolumn quantum energy filter (20 eV slit) |
| Nominal Magnification | 58’180 |
| Pixel Size (Å) | 0.4297 (in super-resolution)<br>0.8594 (for reconstruction) |
| Defocus range (µm) | -0.5 to -1.5 |
| Total electron dose (e/Å$^2$) | 67 |
| Number of movies | ~2000 |
| **Data processing** | |
| Software | RELION 2.1 and RELION 3.0 |
| Initial number of particles | 1’331’906 |
| Final number of particles | 55’245 |

| | |
|---|---|
| Symmetry | C1 |
| Map resolution (Å)<br>FSC threshold | 3.38<br>0.143 |
| Map resolution range (Å) | 3.4 to 3.8 |
| Resolution of sharpened map (Å) | 3.38 |
| Resolution of unsharpened map (Å) | 3.44 |
| Map sharpening B-factor ($Å^2$) | -111 |
| **Refinement** | |
| Refinement package | Phenix (Real-space refinement at 3.40Å) |
| Initial model used (PDB code) | 5YJ6 |
| Map-model CC | |
| CC_mask | 0.73 |
| CC_volume | 0.68 |
| CC_peaks | 0.56 |
| CC_box | 0.62 |
| Model composition | |
| Non-hydrogen atoms | 5180 |

| | |
|---|---|
| Protein residues | 636 |
| Ligands | 0 |
| *B* factors (Å2)(mean) | |
| Protein | 14.14 |
| R.m.s. deviations | |
| Bond lengths (Å) | 0.004 (0) |
| Bond angles (°) | 0.616 (0) |
| Validation | |
| MolProbity score | 1.45 |
| Clashscore | 3.51 |
| Rotamer outliers (%) | 0.56 |
| Ramachandran plot | |
| Favored (%) | 95.58 |
| Allowed (%) | 4.26 |
| Outliers (%) | 0.16 |
| Cb boutliers (%) | 0 |
| CaBLAM outliers (%) | 2.06 |
